# The mitoXplorer 2.0 update: integrating and interpreting mitochondrial expression dynamics within a cellular context

**DOI:** 10.1101/2022.01.31.478461

**Authors:** Fabio Marchiano, Margaux Haering, Bianca Hermine Habermann

## Abstract

Mitochondria are subcellular organelles present in almost all eukaryotic cells, which play a central role in cellular metabolism. Different tissues, health and age conditions are characterised by a difference in mitochondrial structure and composition. The visual data mining platform mitoXplorer 1.0 was developed to explore the expression dynamics of genes associated with mitochondrial functions that could help explain these differences. It however lacked functions aimed at integrating mitochondria in the cellular context and thus, identifying regulators that help mitochondria adapt to cellular needs. To fill this gap, we upgraded the mitoXplorer platform to version 2.0 (mitoXplorer 2.0). In this upgrade we implemented two novel integrative functions, Network Analysis and the transcription factor- (TF-) Enrichment, to specifically help identify signalling or transcriptional regulators of mitochondrial processes. In addition, we implemented several other novel functions to allow the platform to go beyond simple data visualisation, such as an enrichment function for mitochondrial processes, a function to explore time-series data, the possibility to compare datasets across species as well as an IDconverter to help facilitate data upload. We demonstrate the usefulness of these functions in 3 specific use cases. mitoXplorer 2.0 is freely available without login at http://mitoxplorer2.ibdm.univ-mrs.fr.

## INTRODUCTION

Mitochondria are essential organelles in most eukaryotic cells and are involved in a multitude of cellular processes, including cellular energy production, metabolism, signalling or apoptosis. The mitochondrial protein content is estimated to be around 1000 proteins, varying slightly between species (1–3). Most of these proteins (mito-proteins) are encoded in the nucleus and imported into mitochondria in a controlled manner. Only few mito-proteins are encoded in the mitochondrial genome and transcribed and translated within mitochondria, encoding proteins of the electron transport chain (ETC), ribosomal RNAs required (rRNAs) for the mitochondrial ribosome, as well as transfer RNAs (tRNAs). Mitochondria are essential for the functioning of the cell, however the requirements on mitochondria varies with the cell type. Consequently, mitochondrial structure and protein content differ between cell types (4) and we want to understand how mitochondria adapt to their cellular environment.

mitoXplorer version 1 (mitoxplorer 1.0) was developed as a visual data mining platform that allows to analyse the dynamics of gene expression and mutations of all genes with a mitochondrial function (mito-genes) (1). It contains the most complete and up-to-date mitochondrial interactomes for 4 different model species (human, mouse, *Drosophila melanogaster* and budding yeast), as well as 4 different visualisation interfaces that enable users to mine expression and mutation data in a comparative manner, as well as from a single-dataset. mitoXplorer 1.0 was the first platform to mine expression dynamics of mito-genes in this specific manner and was used to identify changes in mito-gene expression dynamics in different conditions. However, it fails to integrate the mitochondrial interactome into the cellular context and thus, to help identify the signalling and transcriptional regulators that adjust mitochondria and their functions to their cellular environment.

We here introduce mitoXplorer version 2 (mitoXplorer 2.0), in which we have added integrative data analysis functions to help address these questions. MitoXplorer 2.0 contains several new functions for data analysis, such an enrichment function for mito-processes, a function to explore time-series data, the possibility to compare datasets across species; as well as two novel integration functions that aim at identifying mitochondrial regulators. First, a function to identify transcriptional regulators of coexpressed mito-genes by making use of our recently published AnnoMiner web-tool (5); and second, a function to identify potential signalling pathways from and to mitochondria, by integrating the mito-interactome in the cellular interactome and subsequently exploring the network neighbourhood of a selected mito-gene based on integrating the network with differential expression data (6). We demonstrate the usefulness of these new functions in three use cases, where we identify potential transcriptional regulators driving the mitochondrial metabolic switch during *Drosophila* flight muscle development (Use Case 1, Figure 2); we identify potential active signalling pathways regulating Ca2+ signalling in Ataxia (Use Case 2, Figure 3); and we explore conserved mito-gene deregulation in human and mouse in Trisomy 21 fibroblasts (Use Case 3, Supplementary Figure S1).

## MATERIAL AND METHODS

Novel functions of mitoXplorer version 2.0, together with those already available in mitoXplorer version 1.0, are shown in Figure 1, as well as Supplementary Figure S2.

**Figure 1:**
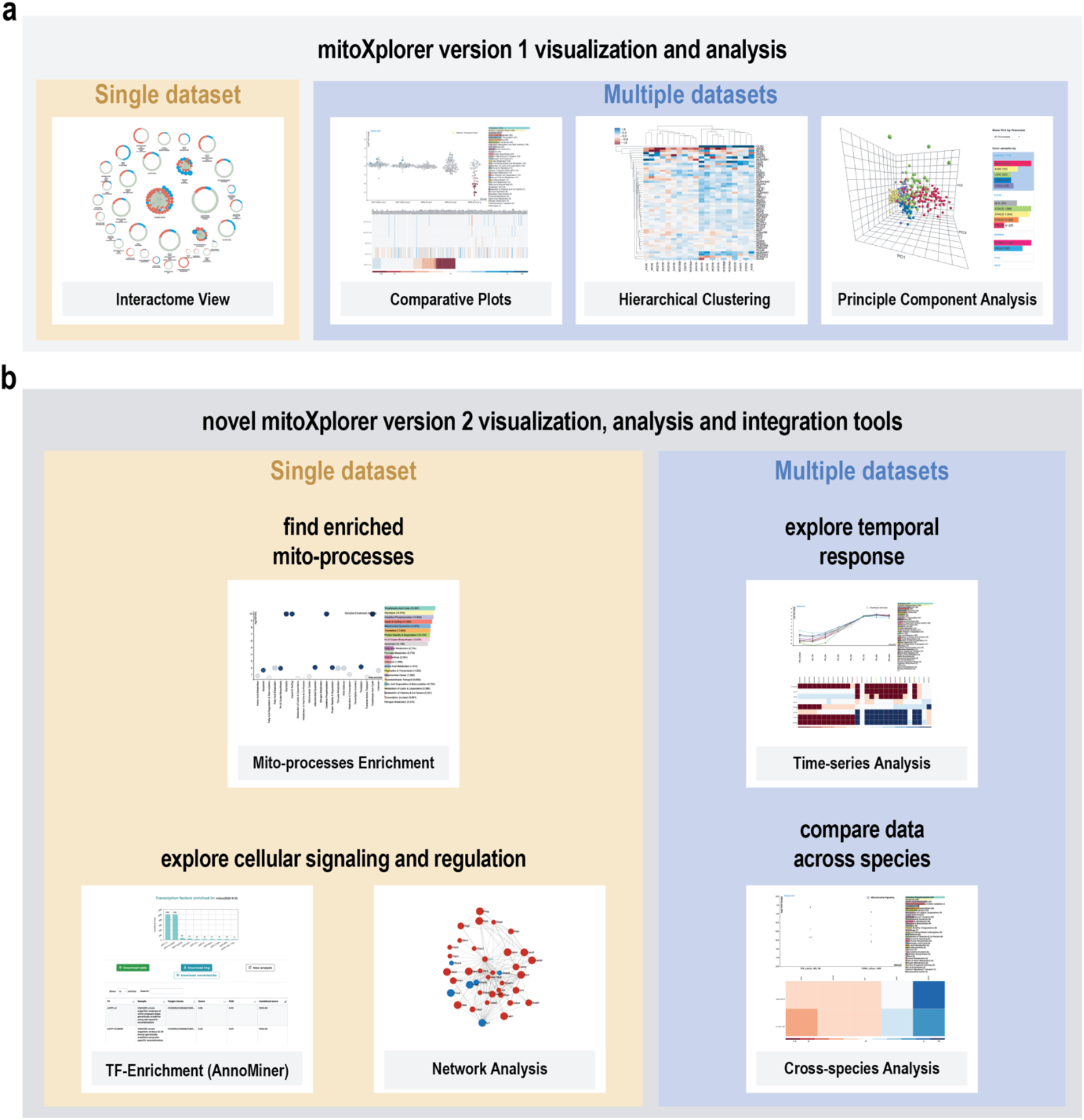
Collection of functions available in mitoXplorer 2.0. **(a)** Visualization and analysis tools available already in mitoXplorer 1.0 include the Interactome View, Comparative Plots, Hierarchical Clustering and Principal Component Analysis. **(b)** Novel functions available in mitoXplorer 2.0 are Enrichment of Mito-Processes, Timeseries Analysis for up to 20 time-points, Cross-species Analysis, as well as the data integration functions of Transcription Factor- (TF-) Enrichment via AnnoMiner, and Network Analysis using the viPEr algorithm.

### Novel functions in mitoXplorer 2.0

#### Mito-process Enrichment Analysis

In order to identify enrichment of our 38 manually curated mito-processes in a dataset, we implemented a Gene Set Enrichment Analysis (GSEA) function. This function helps guide users to focus on important mito-processes of a dataset for subsequent deeper analysis with other functions available in mitoXplorer 2.0.

Mito-process enrichment analysis was coded using the GSEApy python library (7). When p-value and log2FC are both present, genes are ranked by the combined score as defined by Xiao et al. (8):

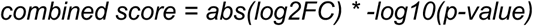

If no p-value is provided, genes are ranked by the absolute value of the log2FC. The gene set enrichment score is tested against a null distribution of enrichment scores generated from 100 permuted gene sets composed of randomly selected genes from the input dataset. The results of the GSEA analysis are shown in the form of an interactive dot-plot, as well as a barplot (see Figure 2 a for an example). In the dot-plot, mito-processes are represented by bubbles, whereby the size reflects the normalised enrichment score (NES) and the colour the combined score. The Y-axis shows the significance of the test (-log10FDR), while the X-axis shows the different mito-processes. By clicking on one bubble representing one mito-process, all values associated with the GSEApy analysis are displayed in the information panel. In the barplot, mito-processes are ranked by combined score, whose value is also given in parenthesis next to the mito-process.

**Figure 2:**
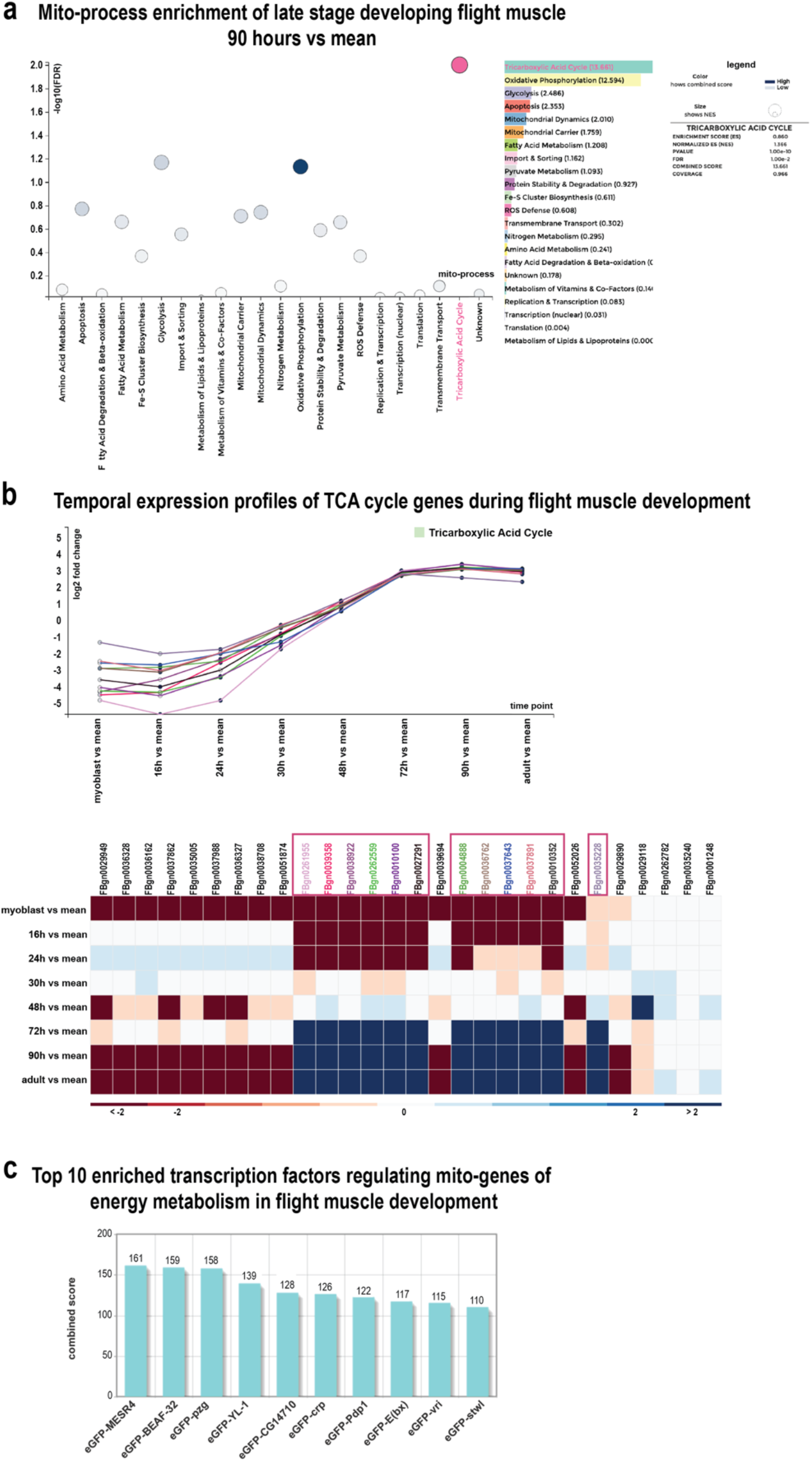
mitoXplorer 2.0 analysis of *Drosophila melanogaster* developing IFM. **(a)** Mito-process Enrichment analysis of 90hrs IFM after puparium formation (APF) revealed an enrichment of TCA, as well as OXPHOS. Mito-processes with a high Combined Score are highlighted in dark blue, the process Tricarboxylic Acid Cycle (TCA) has been selected and appears in pink. **(b)** Time-course Analysis of developing *Drosophila* IFM, showing TCA cycle as an example. Genes that were downregulated early in development (myoblast - 30 hrs APF) and induced in late stages (90 hrs - adult) were selected for display in the scatter plot. Each visualised gene is shown in a different colour. **(c)** Top 10 enriched TFs potentially regulating mito-genes with the described expression pattern (down early, up late) of the mito-processes OXPHOS, OXPHOS mt, and TCA. The plot is directly generated in AnnoMiner upon automatic upload of selected mito-genes within mitoXplorer 2.0’s Time-course Analysis function.

#### Time-course Analysis

In mitoXplorer 1.0, only 6 datasets could be compared to each other with the Correlation Plot. To overcome this limitation and to effectively visualise an increasing number of available time course data, we implemented the Time-course Analysis visualisation. With this function we allow users to visualise up to 20 time points of a temporal dataset. The structure of this visualisation interface is similar to the Comparative Plots, with all its interactivity, except that genes can be individually selected from the sortable heatmap for display in the scatterplot and time points on the X-axis of the scatterplot are connected via a line. The user can thus visualise the temporal gene expression pattern of a single, or multiple mito-genes. Colours are assigned automatically to help visualise individual genes (see Figure 2 b). This visualisation was developed using the d3.js library (9) in order to produce a fully interactive visual interface.

#### Cross-species Analysis

mitoXplorer 2.0 also offers the possibility to compare datasets across species. Cross-species Analysis is possible with the Comparative Plots or the Heatmap function. This will help unravel similarities and understand differences in mitochondrial dynamics between species.

Only the Comparative Plot and the Heatmap are offered for Cross-species Analysis. Orthology information has been included to the backend MySQL database of mitoXplorer by adding orthology ID tables for each model organism. These tables contain dictionaries, having as key the ID of the gene of the model organism of interest and as values the orthologous gene IDs in the other species available in mitoXplorer 2.0. Orthologs between the 4 model species offered by mitoXplorer were initially collected using the last release of BioMart (10). We manually curated these datasets and only 1:1 relations were kept. The user has to first choose ‘Cross species’ from the DATA MINING drop-down menu and subsequently select the species to analyse. In the resulting plots of both visualisation interfaces, the IDs of the first organism selected are used to label genes. However, the user can retrieve the ID of the ortholog of the second organism together with underlying information, by clicking on the respective cell in the heatmap or the bubble in the scatterplot.

#### Integrative analysis functions

In mitoXplorer 2.0, we also wanted to provide the possibility to identify mitochondrial regulators. More specifically, we wanted users to be able to find potential transcriptional regulators of co-regulated mito-genes, as well as identify potentially active signalling pathways from or to mitochondria. The interactivity of mitoXplorer allows one to directly select the gene(s) for both new integrative analysis functions starting from the Comparative Plot, Heatmap or Time-course Analysis functions.

#### TF-Enrichment

Identifying potential transcriptional regulators is done by Transcription factor (TF)-Enrichment analysis by direct re-direction to the AnnoMiner web server (5). In brief, AnnoMiner seeks enriched peaks of transcriptional regulators from ENCODE (11), modENCODE (12) and the modERN (13) databases in the promoter regions of a set of co-regulated genes. This set of genes for an AnnoMiner *TF enrichment analysis* can directly be selected in mitoXplorer 2.0 and uploaded to the AnnoMiner web-server via the TF-Enrichment function (see Supplementary Figure S2 for the TF-Enrichment menu e). The user must then perform TF-Enrichment using the AnnoMiner web server (for a detailed user tutorial of AnnoMiner, see http://chimborazo.ibdm.univ-mrs.fr/AnnoMiner/tutorial.html). As a result, the user receives a list of TFs that potentially regulate the selected gene-set within the AnnoMiner web server environment.

#### Network Analysis

The integrative Network Analysis function aims at identifying signalling paths (active subnetworks) from and to mitochondria, starting from one selected mito-gene. To search for active subnetworks mitoXplorer 2.0 first embeds the mito-interactome into the entire cellular interactome and then allows to integrate differential expression data with the entire interactome. The user can then explore the expression dynamics of the network neighbourhood of a single selected mito-gene and, based on differential expression data, identify potential signalling paths emanating from this gene. To do so, the user has to first activate from the INTEGRATIVE ANALYSIS panel the function Network Analysis (see Supplementary Figure S2 f). Then, he can select the starting mito-gene of interest, the maximum number of steps from the starting point (mito-gene), the log2FC threshold (to define a gene as deregulated or not) and whether unregulated nodes in the results are allowed (see Supplementary Figure S2 for the Network Analysis menu). Results will be displayed directly in a new window in the mitoXplorer platform.

The Network Analysis function has been developed using the NetworkX python library (14). The cellular interactomes were created from the STRING database (v11, (15)) using high-confidence physical protein-protein interactions and have been added to the MySQL database. The network neighbourhood of a selected gene is explored by using the ‘environment search’ algorithm described by the viPEr Cytoscape app (6). Once the analysis has been performed, the active subnetwork is drawn using the d3.js library (9).

#### IDconverter

In order to facilitate data upload, we added an IDconverter to allow usage of any gene identifier. This function converts input gene IDs (eg. ENSEMBL, Entrez, RefSeq, Genecode, Flybase) on the fly to gene symbols, without any intervention from the user. The conversion is performed using the latest release of BioMart (10). BioMart data were downloaded and stored in text format. We retrieved it locally for reasons of speed.

#### Upgrade to Python

In order to enable local deployments and future development of the platform, we upgraded the backend of the web tool from Python 2.7 to Python 3.6.

The new mitoXplorer 2.0 menu is detailed in Supplementary Figure S2 a-f.

### Data processing for use cases 1 and 2

#### Flight muscle data (use case 1)

Temporal expression data of developing *Drosophila melanogaster* indirect flight muscle were taken from (GSE107247, (16)) in the form of raw read counts. In order to obtain differential expression values for the different time-points versus the mean over all time-points, we created two pseudo-replicates of the mean by first calculating the mean over all time-points using one replicate per time-point each. Normalisation of the raw read counts as well as pairwise differential expression analysis were performed using DESeq2.

#### Ataxia data (use case 2)

Data from the Ataxia mouse model ATXN1_82Q_Tg from cerebellum were taken from GSE122099 (17). We downloaded sequencing reads and mapped them to mouse genome version mm10 using STAR (18) with default parameters. FeatureCounts (19) was used to calculate read counts. Normalisation and differential expression analysis was done using DESeq2 via RNfuzzyApp (20). Enrichment analysis was done usine EnrichR (21), for further downstream analysis, we used the KEGG resource (22).

## RESULTS

### Use case 1: Identifying transcriptional regulators of mito-genes during Drosophila melanogaster indirect flight muscle development

We analysed temporal mito-gene expression dynamics during *Drosophila melanogaster* indirect flight muscle (IFM) development using the new mitoXplorer 2.0 functions. In the developing flight muscle, mitochondria undergo significant mitochondrial structural, but also bioenergetic changes (16, 23). To explore these changes further, we used the transcriptomic resource generated by Spletter and colleagues, consisting of transcriptomic data from *Drosophila* isolated IFM at myoblast stage, 16, 24, 30, 48, 72 and 90 hr after puparium formation (APF), as well as the adult (16). To observe temporal changes during IFM development, we compared each time-point to the mean over all time points (see methods) and uploaded the data to mitoXplorer 2.0. We first investigated which mito-processes are enriched during IFM development and used the new Mito-process Enrichment function, using nearly mature IFM at 90 hrs APF. We found mito-processes related to OXPHOS dependent energy metabolism enriched at this time-point (Figure 2 a). We next investigated the temporal expression profiles of mito-genes of the mito-processes OXPHOS, OXPHOS mt (mitochondrial-encoded genes) and Tricarboxylic acid cycle (TCA) using the Time-course Analysis function of mitoXplorer 2.0. As exemplified in Figure 2 b, we found several mito-genes within these 4 mito-processes that were first down-regulated (myoblast stage, 16, 24, 30 hrs APF) and subsequently strongly induced from 72 hrs to adult stage. We wanted to identify potential transcriptional regulators of the co-regulated genes from these 3 processes, using mitoXplorer’s integrative analysis function TF-Enrichment. We therefore selected all genes from the 4 processes that showed the above described temporal changes and uploaded them via mitoXplorer’s TF-Enrichment function to AnnoMiner (see Supplementary Table S1 a for the list of genes selected). AnnoMiner returned several interesting TFs as enriched in the promoters of the co-regulated mito-genes (Supplementary Table S1 b). Among the top 10 hits are MESR4, which has been shown to be involved in development and in the cellular response to hypoxia (24); pzg, which is known to regulate developmental processes (25, 26); Pdp1, which is known to regulate muscle genes (27); crp, which controls cell growth and tracheal terminal branching (28); or vri, which has been shown to be involved in tracheal development (29, 30).

To conclude, this use case demonstrates how the new functions of mitoXplorer 2.0, Mito-process Enrichment, Time-course Analysis as well as the integrative analysis function TF-Enrichment could help identify the gene expression dynamics behind the mitochondrial bioenergetic switch in developing IFM and predict potential transcriptional regulators responsible for this switch.

### Use Case 2: Identification of signalling cascades regulating Ca2+ signalling in a mouse model of Spinocerebellar Ataxia 1 type 1 (SCA1)

Spinocerebellar Ataxias (SCAs) is a group of dominantly inherited neurodegenerative diseases, which is defined by a loss of coordination of body movements and cerebellar degeneration. They can be caused by mutations in nearly 40 genes and are currently untreatable. A disruption of Ca2+ signalling in cerebellar cells, and more specifically in the Purkinje neurons, is considered to play a key role in disease onset and progression (31). We wanted to explore the contribution of mitochondria to Ca2+ signalling deregulation in SCAs. To this end, we used a mouse model of Spinocerebellar ataxia type 1 (SCA1). SCA1 affects the cerebellum, as well as the inferior olive and is caused by poly-glutamine (polyQ) expansion in the ATAXIN1 gene, which encodes a transcriptional regulator (32). We used RNA expression data from a mouse model of SCA1 from (17), who looked at cerebellum and inferior olive of 5 and 12 week old transgenic ATXN1_82Q (ATXN1_82Q Tg) mice. We focused mitoXplorer analysis on the cerebellum and were mostly interested in the temporal differences in potential pathways affecting mitochondrial Ca2+ signalling and transport.

We uploaded differential expression data of ATXN1_82Q Tg compared against wild-type (WT) at 5 and 12 weeks, as well as the temporal comparisons of WT (5 to 12 weeks), and ATXN1_82Q Tg (5 to 12 weeks) to mitoXplorer 2.0 and used the Comparative Plots to visualise changes in ‘Ca2+ Signalling and Transport’. Among the genes most affected in this mito-process was Itpr1 (Figure 3 a), an intracellular receptor for inositol 1,4,5-trisphosphate that is located to the endoplasmic reticulum (ER), is one of the main regulators of mitochondrial Ca2+ signalling (33) and is itself regulated by mitochondria (34). Furthermore, mutations in ITPR1 cause Spinocerebellar Ataxia Type 15 (SCA 15, (35)). We next explored the network neighbourhood of Itpr1 at 5 and 12 weeks using the integrative Network Analysis function (Figure 3 b, c; no deregulated nodes allowed, log2FC >= |1.2|, step-size 3). The extracted active subnetwork surrounding Itpr1 is substantially larger at 12 weeks with more differentially expressed nodes within a 3-step neighbourhood (Figure 3 c) compared to the earlier stage (Figure 3 b). Many of the enriched pathways overlap between the two time-points, such as Calcium signalling, Neuroactive ligand-receptor interaction, or Phospatidylinositol signalling. At 12 weeks, some more metabolic pathways, as well as Spinocerebellar ataxia appear as being enriched (Supplementary Tables 2 b-d). Moreover, 3 genes responsible for SCAs are found in the 12 weeks network, including Itpr1 itself together with Trpc3 and Cacna1g (Supplementary Table S2 e).

**Figure 3:**
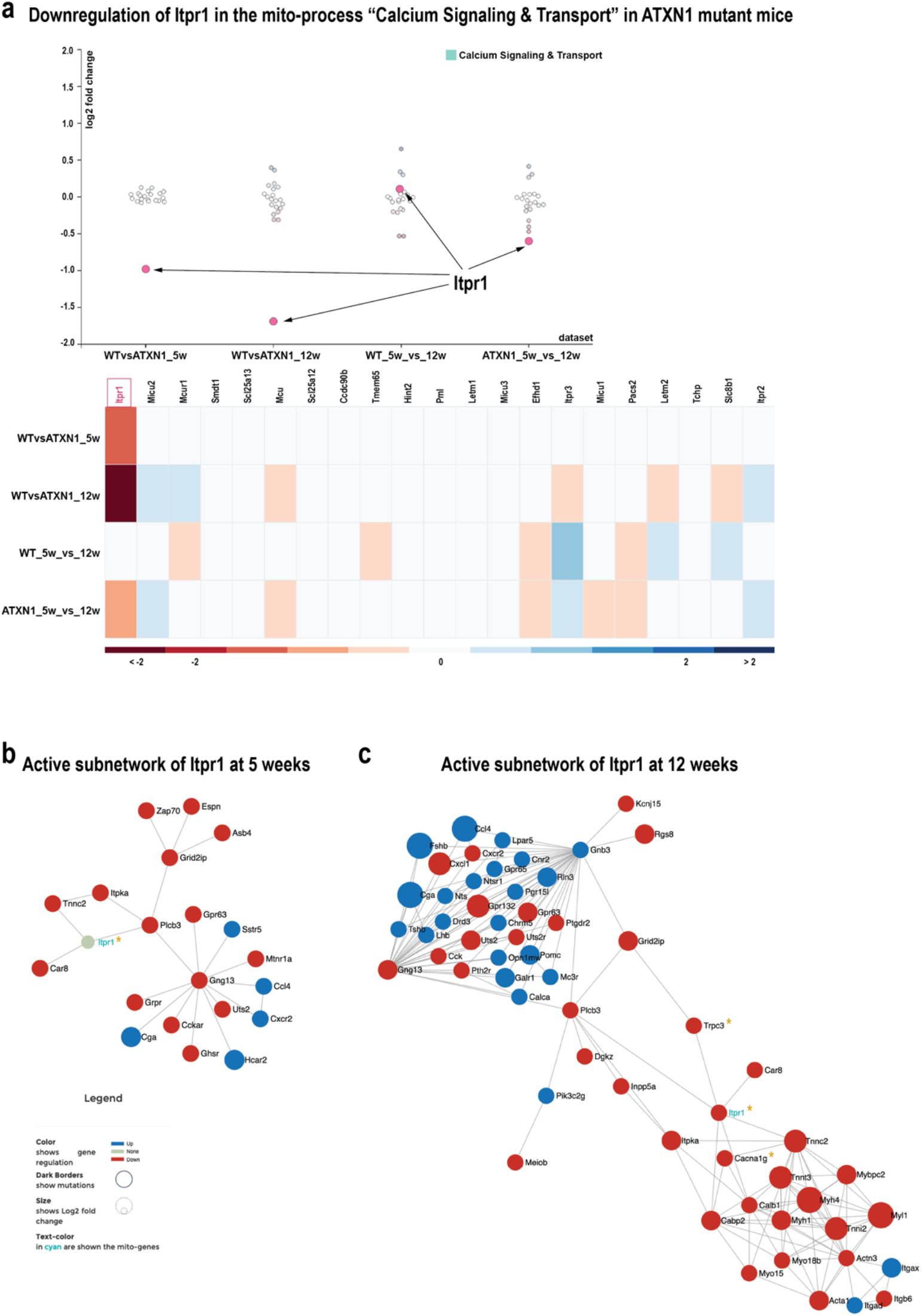
Ca2+ signalling and transport is affected in spinocerebellar ataxia type 1 (SCA1). **(a)** One of the most affected genes in the ATXN1_Q82 transgene mouse model for SCA1 is Itpr1, which localises to the ER and regulates mitochondrial Ca2+ signalling. **(b)** Network neighbourhood of Itpr1 in 5-week-old ATXN1_Q82 Tg mice. **(c)** Network neighbourhood of Itpr1 at 12 weeks’ time. Genes found to cause a form of SCA are marked with an asterisk. Mito-genes are labelled in cyan while non-mitochondrial genes are labelled in black. The user can interactively change genes disposition and labels in order to show different information (gene name or ID, mutation, log2FC). The visualisation is information dense. Node size, colour and border represent magnitude of the log2FC, direction of the deregulation (up or down) and mutation information, when available.

In conclusion, we used the Comparative Plot and Network Analysis functions to investigate Ca2+ signalling de-regulation in a SCA1 mouse model and found several signalling pathways consistently induced from 5 to 12 weeks in the cerebellum. Moreover, two more genes related to SCA were found in the 12 weeks active subnetwork.

## DISCUSSION

Here we present an important upgrade of the mitoXplorer visual data mining platform. While mitoXplorer version 1.0 (1) allowed to inspect mitochondrial datasets in multiple ways, mitoXplorer 2.0’s integrative analysis functions permit the user to go beyond simple data visualisation. Our new integrative functions can be used to better understand the interaction between mitochondria and their cellular environment, as we have shown in two of our use cases. In the first use case we showed how the new functions of mitoXplorer 2.0 the Mito-process Enrichment, Time-course Analysis as well as TF-Enrichment can be used to identify key transcriptional regulators of important mitochondrial processes. In the second one, we employed Network Analysis which, by integrating the mitochondrial interactome into the complete cellular interactome, allows the identification of important non-mitochondrial genes that are connected with and potentially regulate mitochondrial processes. These results demonstrate the power of mitoXplorer 2.0’s downstream and integrative analysis functions. For TF-Enrichment, we decided to rely on AnnoMiner (5). By connecting mitoXplorer 2.0 to AnnoMiner, direct data transfer from one to the other is possible to execute AnnoMiner’s *TF enrichment analysis,* to identify potential transcriptional regulators of mitochondrial functions. To date several tools for TF-Enrichment are available but to our knowledge only few of these, namely modEnrichr (36), iCisTarget (37) and AnnoMiner, work on model organisms other than human and mouse and thus can be used for three out of the 4 model organisms present in mitoXplorer 2.0: human, mouse, and *D. melanogaster.* AnnoMiner has also data for *Caenorhabditis elegans,* a model species we plan to offer soon on mitoXplorer 2.0. For Network Analysis, we implemented a function for active subnetwork extraction starting from a single gene based on the algorithm of the Cytoscape app viPEr (6). The exploration of the network neighbourhood of a gene using viPEr requires as user input only the number of steps from one selected mito-gene of interest together with a log2FC threshold to define genes as deregulated. The algorithm thus avoids cumbersome tuning of parameters. The fact that we implemented this algorithm using NetworkX (14) and D3.js (9) permits a fully interactive exploration of the extracted subnetwork, facilitating its interpretation. In the future, we plan to include downstream analysis of the resulting network, such as pathway annotation through enrichment analysis. This should help to better identify potential signalling paths regulating mitochondrial functions. We find these two integrative analysis functions as an important novel upgrade of mitoXplorer, which should help to unravel how mito-gene expression and thus, mitochondrial functions within the cell and thus understand, how mitochondria are able to adjust in content and function to a specific cellular environment.

Additional future developments we currently work on include mitochondrial metabolic modelling using Flux Balance Analysis (FBA), as well as logical modelling. Combining mitoXplorer directly with metabolic modelling would allow users to assess the metabolic states of biological samples based one mito-gene expression data directly within the mitoXplorer platform. Starting mitochondrial models are available for human and budding yeast (38–40), which we will adapt to our needs. In addition, we plan to extend mitoXplorer to allow analysis of single-cell RNA-seq data of tissues on cell-type level. All these further implementations are simplified by the modular fashion in which mitoXplorer 2.0 has been developed.

## AVAILABILITY

mitoXplorer 2.0 is available as a web-server at: http://mitoxplorer2.ibdm.univ-mrs.fr. The source code together with installation instructions are available in our GitLab repository (https://gitlab.com/Fabio_M/MitoX2).

## SUPPLEMENTARY DATA

**Use case 3 and Supplementary Figure S1:** Comparison of differential expression in ROS defense from fibroblasts of human Trisomy 21 with fibroblasts from a mouse model for Trisomy 21.

**Supplementary Figure S2: New mitoXplorer 2.0 menu.** Detailed description of the new menu of mitoXplorer 2.0.

**Supplementary Table S1: (a)** *Drosophila* gene lists uploaded to AnnoMiner. **(b)** Enriched TFs identified for **(a)** by AnnoMiner.

**Supplementary Table S2: (a)** Gene lists of 5- and 12-weeks active subnetworks of the ATXN1_82Q Tg mouse model. **(b)** Enriched KEGG pathways of 5 weeks ATXN1_82Q Tg gene list. **(c)** Enriched KEGG pathways of 12 weeks ATXN1_82Q Tg gene list. **(d)** KEGG pathways with >3 genes from the 5 week and 12-week networks. **(e)** Genes related to Spinocerebellar Ataxias in the 5 week and 12-week network.

## AUTHOR CONTRIBUTIONS

The upgrade of the mitoXplorer web-server was developed by F.M. Data analysis was done by B.H.H., F.M. and M.H. The project was conceived by B.H.H., the manuscript was written by F.M. and B.H.H with input from M.H. All authors read and approved the final version of the manuscript.

## ACKNOWLEDGEMENT

We thank Alice Carrier, Benoit Giannesini, Frank Schnorrer, Nuno Luis, Jerome Avellaneda, as well as the members of the IBDM Computational Biology team for helpful discussion on mitoXplorer 2.0. We thank Stephen Chapman for critical reading of the manuscript.

## FUNDING

This work was supported by the Agence National de Recherche (ANR) grant number ANR-18-CE45-0016-01 MITO-DYNAMICS and Fondation pour la Recherche Médicale (FRM) grant AtaxiaXplorer (MND202003011460) awarded to BHH, the Centre National de la Recherche Scientifique (CNRS), Aix-Marseille University and the IBDM UMR 7288.

## CONFLICT OF INTEREST

The authors do not have any conflict of interest.

## Notes

http://mitoxplorer2.ibdm.univ-mrs.fr

https://gitlab.com/Fabio_M/MitoX2

### Competing Interest Statement

The authors have declared no competing interest.

