## Supplementary Data, Supplementary Figure S1, Supplementary Figure S2 for "The mitoXplorer 2.0 update: integrating and interpreting mitochondrial expression dynamics within a cellular context"

**Use case 3 and Supplementary Figure S1:** Comparison of differential expression in ROS defense from fibroblasts of human Trisomy 21 with fibroblasts from a mouse model for Trisomy 21.

**Supplementary Figure S2: New mitoXplorer 2.0 menu.** Detailed description of the new menu of mitoXplorer 2.0.

**Supplementary Table S1: (a)** *Drosophila* gene lists uploaded to AnnoMiner. **(b)** Enriched TFs identified for **(a)** by AnnoMiner.

**Supplementary Table S2: (a)** Gene lists of 5- and 12-weeks active subnetworks of the ATXN1_82Q Tg mouse model. **(b)** Enriched KEGG pathways of 5 weeks ATXN1_82Q Tg gene list. **(c)** Enriched KEGG pathways of 12 weeks ATXN1_82Q Tg gene list. **(d)** KEGG pathways with >3 genes from the 5 week and 12-week networks. **(e)** Genes related to Spinocerebellar Ataxias in the 5 week and 12-week network**.**

##### Use case 3: Comparison of differential expression in ROS defense from fibroblasts of human Trisomy 21 with fibroblasts from a mouse model for Trisomy 21

We wanted to know, whether differential expression patterns of mito-genes are similar in fibroblasts from human monozygotic twins discordant for Trisomy 21 model and a mouse model of Trisomy 21. We used data from (1), which were already uploaded to mitoXplorer and invoked the Cross-species function of mitoXplorer 2.0. We used the comparative Plots to compare the two datasets of trisomic fibroblasts, one from human monozygotic twins discordant for Trisomy 21, and the other from fibroblasts of a mouse model of Trisomy 21, Ts65DN. In the process ROS defense, we find that most genes are co-regulated between both species, except for the Neuroglobin (NGB), a globin-like gene that protects against neurodegeneration, hypoxia, ischemia or nutrient deprivation (2), which is strongly induced in murine trisomic fibroblasts.

#### Supplementary Figure S1

**Cross-species comparison of human Trisomy 21 fibroblasts against fibroblast from a Trisomy 21 mouse model (Ts65DN)**

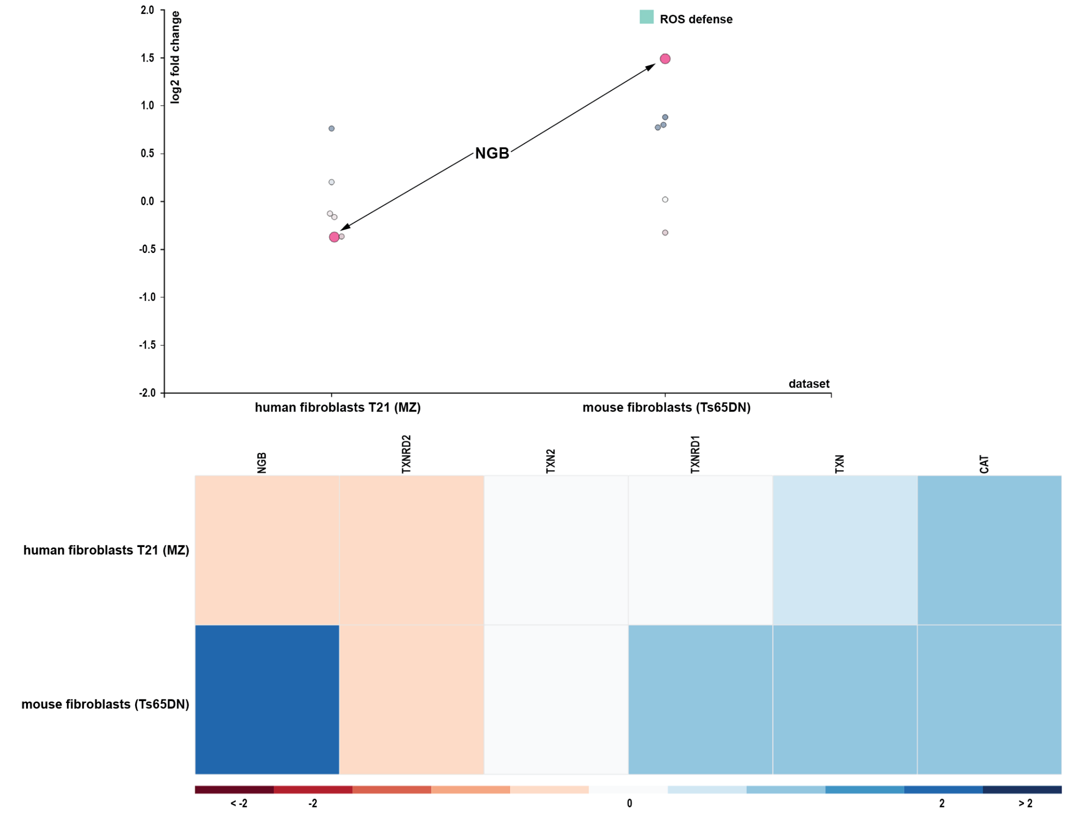


**Figure S1: Cross-species comparison of differential expression data between data from human fibroblasts from monozygotic twins discordant for Trisomy 21 and fibroblasts from a mouse model for Trisomy 21 (Ts65DN).** Differential expression data from human and mouse Trisomy 21 fibroblasts were compared against their respective wild-type controls (data available in mitoXplorer (3). Red color indicates downregulation, blue color up-regulation. The gene NGB has been selected in the scatterplot.

### Supplementary Figure S2

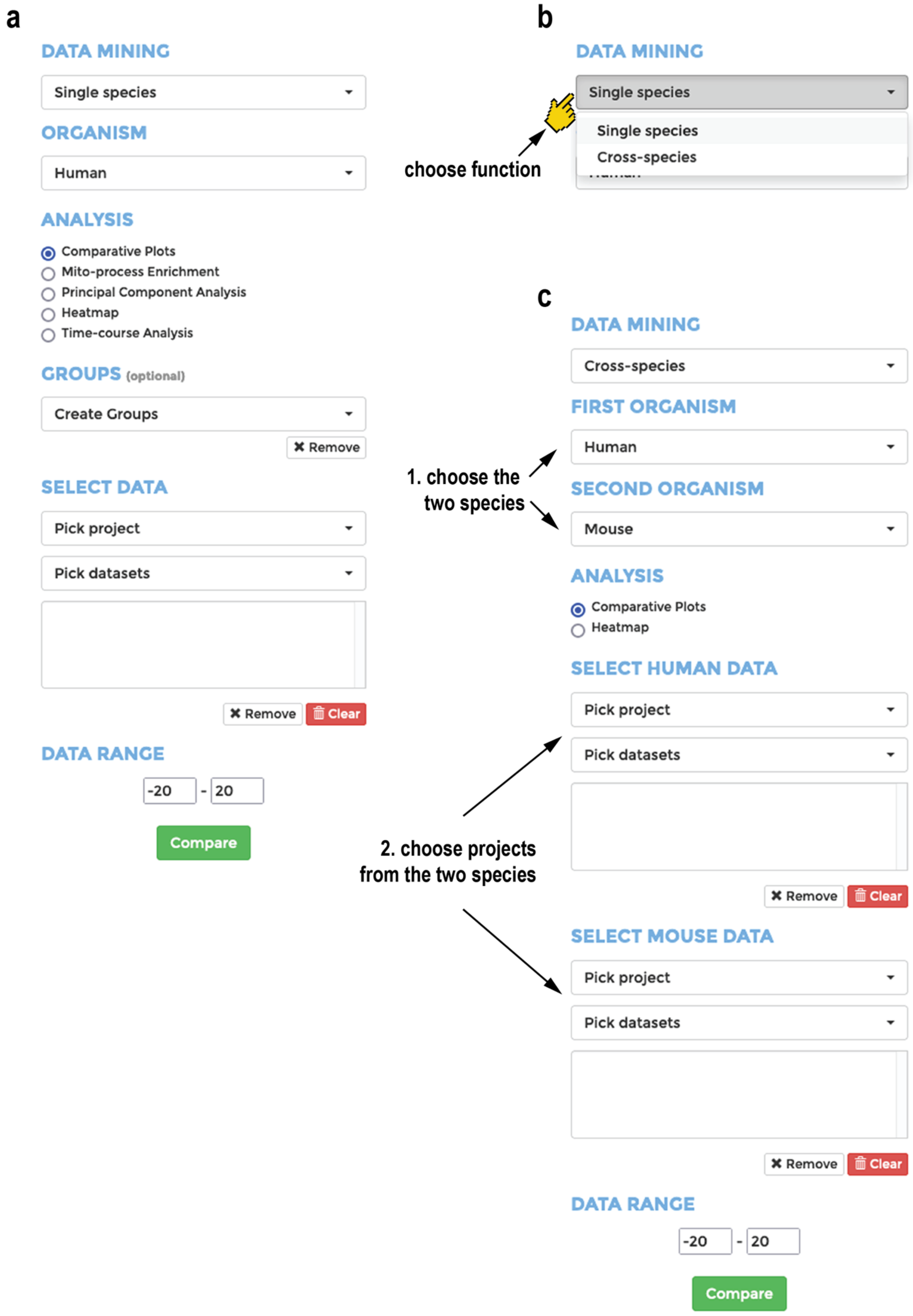

#### Supplementary Figure S2

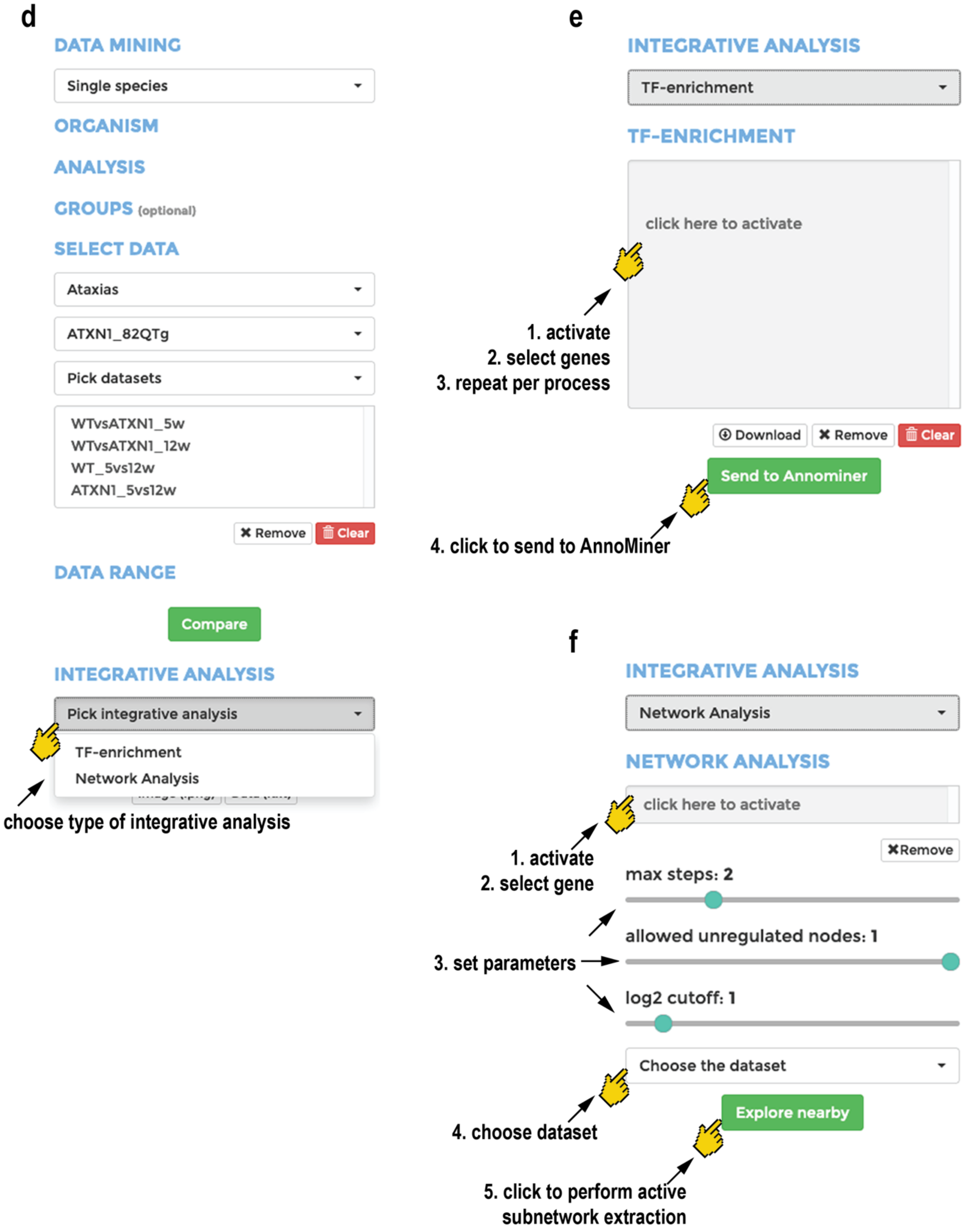


**Supplementary Figure S2: New menu of mitoXplorer 2.0. (a)** New menu functions of mitoXplorer main menu includes the DATA MINING panel for choosing between single species and cross-species; New functions in the ANALYSIS panel include Mito-process Enrichment and Time-Course Analysis. **(b)** When clicking on the drop-down menu in DATA MINING, the user can choose between Single species and Cross-species. **(c)** When choosing Cross-species analysis, 1. the two organisms have to be chosen; and 2. the datasets to be compared from both organisms have to be chosen. As analysis functions, Comparative Plots as well as Heatmaps are offered **(d)** Once a dataset has been uploaded, the INTEGRATIVE ANALYSIS panel appears. From here, the user can choose in a drop-down menu TF-enrichment and Network Analysis **(e)** When choosing TF-enrichment, a window appears that needs to be clicked (1.), before genes can be selected (2.). When genes from a second mito-process should be added, the window needs to be clicked again (3.). Once the user is satisfied with the selected gene list, it can be sent to AnnoMiner by clicking on the green box called ‘Send to AnnoMiner’ (4.). The gene list can also be downloaded when clicking on the box called ‘Download’. The user will be directed to AnnoMiner to look for enriched TFs (for details on the usage of AnnoMiner, see (4) and also the video instructions provided on the AnnoMiner web-site (http://chimborazo.ibdm.univ-mrs.fr/AnnoMiner/tutorial.html). **(f)** When Network Analysis is chosen as downstream analysis, several items appear: a box that must be activated (1.) and where a single gene can be selected (2.); a set of parameters (3.), which include the maximal step size of the resulting active subnetwork, the number of nodes in the paths allowed without being differentially expressed; and the log2FC cutoff that defines differentially expressed genes; a drop-down menu from which a dataset to be used for the active subnetwork extraction has to be chosen (4.); finally, by clicking on the green box called ‘Explore nearby’, the gene together with the dataset and chosen parameters will be sent to the subnetwork extraction function. A resulting network will appear in a separate browser window within the mitoXplorer platform.
